# A multivariate genome-wide association analysis of 10 LDL subfractions, and their response to statin treatment, in 1868 Caucasians

**DOI:** 10.1101/011270

**Authors:** Heejung Shim, Daniel I. Chasman, Joshua D. Smith, Samia Mora, Paul M. Ridker, Deborah A. Nickerson, Ronald M. Krauss, Matthew Stephens

## Abstract

We conducted a genome-wide association analysis of 7 subfractions of low density lipoproteins (LDLs) and 3 subfractions of intermediate density lipoproteins (IDLs) measured by gradient gel electrophoresis, and their response to statin treatment, in 1868 individuals of European ancestry from the Pharmacogenomics and Risk of Cardiovascular Disease study. Our analyses identified four previously-implicated loci (SORT1, APOE, LPA, and CETP) as containing variants that are very strongly associated with lipoprotein subfractions (log_10_Bayes Factor > 15). Subsequent conditional analyses suggest that three of these (APOE, LPA and CETP) likely harbor multiple independently associated SNPs. Further, while different variants typically showed different characteristic patterns of association with combinations of subfractions, the two SNPs in CETP show strikingly similar patterns - both in our original data and in a replication cohort - consistent with a common underlying molecular mechanism. Notably, the CETP variants are very strongly associated with LDL subfractions, despite showing no association with total LDLs in our study, illustrating the potential value of the more detailed phenotypic measurements. In contrast with these strong subfraction associations, genetic association analysis of subfraction response to statins showed much weaker signals (none exceeding log_10_ Bayes Factor of 6). However, two SNPs (in APOE and LPA) previously-reported to be associated with LDL statin response do show some modest evidence for association in our data, and the subfraction response profiles at the LPA SNP are consistent with the LPA association, with response likely being due primarily to resistance of Lp(a) particles to statin therapy. An additional important feature of our analysis is that, unlike most previous analyses of multiple related phenotypes, we analyzed the subfractions jointly, rather than one at a time. Comparisons of our multivariate analyses with standard univariate analyses demonstrate that multivariate analyses can substantially increase power to detect associations. Software implementing our multivariate analysis methods is available at http://stephenslab.uchicago.edu/software.html

**Author Summary:** Levels of plasma lipids and lipoproteins are related to risk of cardiovascular disease (CVD), and because of this, considerable attention has been devoted to genetic association analyses of lipid-related measures. In addition, motivated by the fact that statins are widely prescribed to lower plasma low density lipoprotein (LDL) cholesterol and CVD risk, and that response to statins has a genetic component, several studies have searched for genetic associations with response of lipid related phenotypes to statin treatment. Here, in 1868 individuals of European ancestry from the Pharmacogenomics and Risk of Cardiovascular Disease study, we have conducted genetic association analyses of 7 subfractions of LDLs and 3 subfractions of intermediate density lipoproteins (IDLs) measured by gradient gel electrophoresis, and their response to statin treatment. These phenotypic measurements offer higher resolution information on LDLs and IDLs than available previously. Therefore, our study provides a more detailed picture of association with the entire IDL/LDL subfraction profile than any prior genetic association studies of either lipid-related measures or their response to statin treatment. Moreover, unlike most previous analyses of multiple related measurements, we analyzed the subfractions jointly, rather than one at a time. Our results demonstrate that joint analyses of related measurements can considerably increase power to detect associations compared with conventional univariate analyses.

## Introduction

Levels of plasma lipids and lipoproteins are related to risk of cardiovascular disease, and because of this, considerable attention has been devoted to genetic association analyses of lipid-related measures. The largest of these studies are genetic association analyses for plasma concentrations of the common clinical lipid phentoypes: total cholesterol (TC), LDL-cholesterol (LDL-C), HDL-cholesterol (HDL-C), and triglycerides (TG). For example, [1] performed a meta-analysis of these traits in > 100, 000 individuals of European ancestry, and identified a total of 95 associated loci. Other (smaller, although still substantial) studies considered genetic associations with more detailed lipid-related measurements, specifically plasma concentrations of subfractions of very low density lipoproteins (VLDLs), intermediate density lipoproteins (IDLs), LDLs, and HDLs, measured by NMR [2–4]. And, motivated by the fact that statin drugs are widely used to treat lipid phenotypes, and that response to these drugs has a genetic component, several studies have searched for genetic associations with response of LDL-C, HDL-C, TC and TG to statin treatment [5–7].

In this paper we describe genetic association analyses of 7 subfractions of LDLs and 3 subfractions of IDLs measured by gradient gel electrophoresis, and of their response to statin therapy. Although our sample size is smaller than many recent lipid association studies (1868 individuals), our phenotypic measurements provide higher resolution information for IDLs and LDLs than prior genetic association studies of either statin-treated or untreated samples. Specifically, our 10 subfractions of IDLs and LDLs compare with 3-4 size subfractions in the NMR-based studies, and no previous genome-wide association study of lipoprotein response to statin therapy has considered subfraction data. While our smaller sample size limits what we can say about genetic variants with small effects, for variants with sufficiently strong associations our data provide a more detailed picture of their associations with the entire IDL/LDL subfraction profile than any previous study. We find several examples of SNPs that are only very weakly associated with total LDL-C in our study, but are much more strongly associated with one or more individual subfractions. Of particular note, we highlight two independently-associated variants in the CETP gene that have no overall effect on total LDL-C in our study, but a strong effect on several individual IDL/LDL subfractions (in addition to a well-established strong effect on total HDL-C).

In addition to the detailed nature of the phenotypes, our study also differs from most previous studies in our use of *multivariate* association analysis of related phenotypes, rather than treating each phenotype separately. Our results illustrate that association analysis of multivariate phenotypes can substantially increase the strength of association signals compared with conventional univariate analyses.

## Methods

### Study populations and genotype data

All samples in our analysis were derived from the Pharmacogenomics and Risk of Cardiovascular Disease (PARC) study. The study population, experimental design, and genotyping procedures have been described in detail previously [6]. Briefly, this study contains individuals from two statin trials: the Cholesterol and Pharmacogenetics (CAP) study [8], and the Pravastatin Inflammation/CRP Evaluation (PRINCE) study [9]. The PRINCE study consists of two cohorts, one containing individuals with history of CVD (secondary prevention cohort) and the other containing individuals with no history of CVD (primary prevention cohort). Participant characteristics are summarized in Table 1.

**Table 1.**
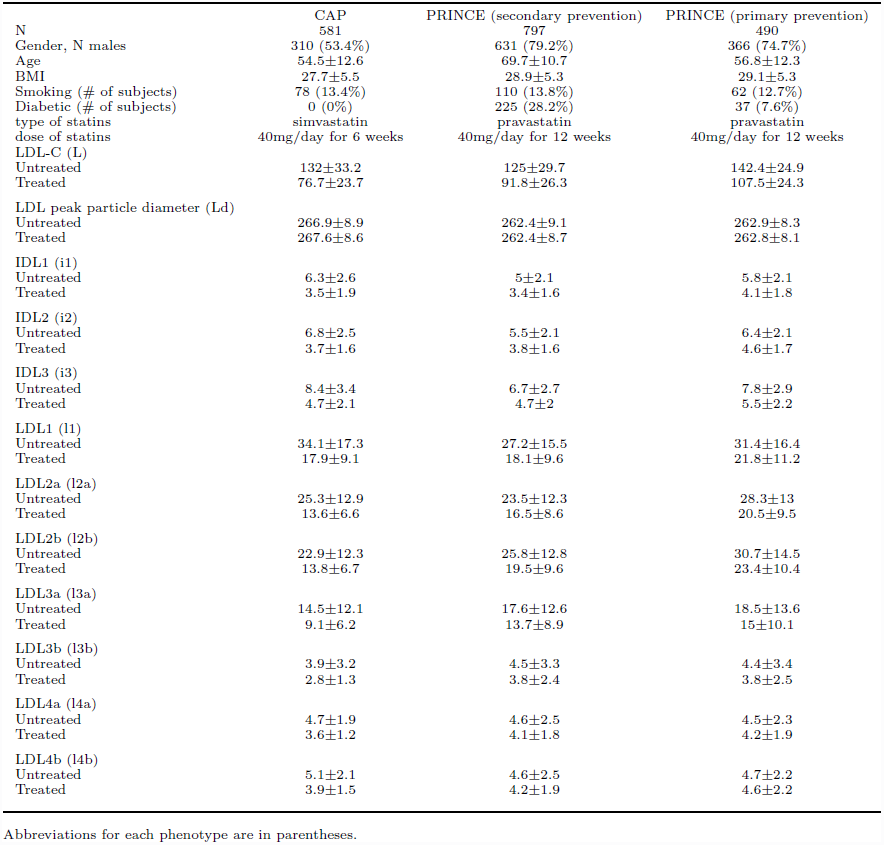
Summary of phenotype distributions (meanstandard deviation) for individuals in our analysis, stratified by study/cohort.

Genotyping was conducted in two stages. The first stage individuals were genotyped on the Illumina HumanHap300 bead chip and the second stage individuals were genotyped on the Illumina HumanQuad610 bead chip and a custom-made iSelect chip. The HumanHap300 and the HumanQuad610 chips (henceforth referred to as the 300K chip and the 610K chip) were designed to tag common variation among individuals of European ancestry while 12,959 SNPs in the iSelect chip were selected to increase coverage of candidate SNPs for cardiovascular disease regardless of minor allele frequency (MAF). Our analyses reported here utilized a total of 1,868 Caucasian individuals for whom complete LDL subfraction phenotype data were available (see below).

To maximize genomic coverage and combine the multiple groups genotyped on different SNP chips, we performed genotype imputation [10] [11], using an imputation protocol that has been previously described [12]. Briefly, genotype imputation was performed using IMPUTE2 [10] with an integrated reference panel that included 120 CEU haplotypes from the 1000 Genomes Pilot Project (“1000G”) [13] and 1910 worldwide haplotypes from the HapMap Phase 3 Project (“HM3”) [14]. This procedure generated genotypes (either genotyped or imputed) for 7,836,525 SNPs.

### Phenotypic measurements and normalisation

LDL cholesterol was estimated by the formula of [15] using measured total cholesterol, HDL cholesterol, and triglyceride. All lipid measurements were consistently in range as determined by the NHLBI-CDC Lipid Standardization Program. Gradient gel electrophoresis estimates of LDL subclass concentrations and LDL peak diameter were determined from whole plasma by 2%-14% non-denaturing polyacrylamide gradient gel electrophoresis as described previously [16] except that Sudan Black was used as the lipid stain [17], and the size range was extended to include IDL [18] since the Friedewald formula provides a measure of cholesterol in the density range that includes IDL as well as LDL particles. The subfractions analyzed and their corresponding particle size intervals were: LDL4b (22.0 - 23.2 nm), LDL4a (23.3 - 24.1 nm), LDL3b (24.2 - 24.6 nm), LDL3a (24.7 - 25.5 nm), LDL2b (25.6 - 26.4 nm), LDL2a (26.5 - 27.1 nm), LDL1 (27.2 - 28.5 nm), IDL3 (28.6 - 29.0 nm), IDL2 (29.1 - 29.6 nm) and IDL1 (29.7 - 30.3nm). The cholesterol concentrations of these subfractions were determined by multiplying percent of the total stained LDL and IDL for each subfraction by the LDL cholesterol value [17]. In both studies, the LDL subfractions and LDL cholesterol values were estimated from blood samples taken on one visit before statin treatment and one visit during the treatment (after 6 weeks in CAP and 12 weeks in PRINCE).

Following [6] we separately normalized the 12 LDL-related phenotypes (total LDL-C, LDL peak diameter, and the 10 LDL subfractions), by the following normalization procedure, which aims to reduce the influence of outlying observations and avoid false positive association due to systematic differences between studies/cohorts. We partitioned the individuals into six strata, corresponding to the two stages of genotyping in each of three groups (CAP, PRINCE primary prevention and PRINCE secondary prevention). Within each stratum we separately quantile transformed the pre-treatment and post-treatment measurements to a standard normal distribution to yield pre-treatment and post-treatment phenotypes, *P* and *T*. Then we calculated both the average *A* = (*T* + *P*)/2, and the difference *D* = (*T -P*) of the transformed measurements for each individual. (The motivation is that analyses of *A* are well-powered to identify associations that occur in both *T* and *P*, whereas analyses of *D* are aimed at identifying associations with statin response.) We then separately corrected *A* and *D* for covariates (age, log(BMI), sex, and smoking status) within each stratum using a standard multiple linear regression, to yield covariate-corrected phenotypes *A′* and *D′* (obtained from the residuals of the multiple regression). Finally, each of *A′* and *D′* were again quantile transformed to a standard normal distribution within each strata, to yield final phenotypes *Ã* and 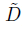. Since previous analyses of the PARC study suggest little potential for spurious associations due to cryptic population stratification within strata [6], we did not further control for population substructure within strata.

### Replication cohort, genotype data, phenotypic measurements and normalization

Replication of our CETP associations was performed among individuals of European descent from the JUPITER study [7,19]. Specifically, we examined genotype data at our two replication SNPs, and phenotype data among up to 6745 JUPITER participants on VLDL, IDL and LDL subfractions measurements determined by an ion mobility assay [20, 21]. We focused on the 12 ion-mobility phenotypes that correspond most closely to the 12 phenotypes in our study, namely: total LDL (L), LDL peak diameter (Ld), VLDL small (i1), IDL1 (i2), IDL2 (i3), LDL1 (l1), LDL2a (l2a), LDL medium small (l2b), LDL small (l3a), LDL3b (l3b), LDL4a (l4a), LDL4b (l4b).

Before association analysis each phenotype was quantile transformed to a standard normal distribution, and then adjusted for covariates (age, sex, and region: North America vs Europe) using a standard multiple linear regression. The covariate-corrected phenotypes (i.e. the residuals of the multiple regression) were used in replication analyses. For the top SNP we regressed these covariate-corrected phenotypes against SNP genotype to estimate an effect for each phenotype. We then took the residuals from this regression (to control for the effect of the top SNP) and regressed them against SNP genotype at the secondary SNP to estimate remaining effects at the secondary SNP.

### Multivariate association analyses

We performed multivariate association analyses using methods described in detail in [22]. To outline these methods, consider analyzing *d* phenotypes *Y*_*1*_,…, *Y*_*d*_, and assessing their association with a single SNP, whose genotypes are denoted by *g*. Instead of performing a single test of association, the idea is to compare different models for the way that *Y*_1_,…, *Y*_*d*_ could be associated with *g*. The null model is that none of them are associated with *g*. Then there are a large number of alternative models, in which different subsets of the variables are associated with *g*. For example, one alternative is that *Y*_1_ alone is associated with *g*, and the others are unassociated. Another is that *Y*_2_ alone is associated with *g*. Another is that all of them are associated with *g*, etc. In addition, the method allows that some phenotypes might be “indirectly” associated with *g*, which means that their association with *g* is mediated entirely through the directly associated phenotypes. (Specifically, these phenotype are associated with *g*, but are conditionally independent of *g* given the directly associated phenotypes.)

Formally, consider partitioning the phenotypes into three groups, *U*, *D* and *I* where *U* denotes the phenotypes that are unassociated with *g*, *D* denotes the phenotypes that are “directly” associated with *g*, and *I* denotes the phenotypes that are “indirectly” associated with *g*. The support in the data for each possible partition *γ* = (*U, D, I*), compared with the null (*γ* = *γ*_0_, where *γ*_0_ is the partition with all phenotypes in *U*), is given by the Bayes Factor BF_*γ*_ = P(Y|*g,γ*)/P(Y|*γ*_0_). Here we use expressions for these Bayes Factors from a multivariate normal model for *Y* [22]. Since in practice any of the alternative hypotheses may hold, the overall evidence against the null hypothesis is given by a weighted average of these Bayes Factors

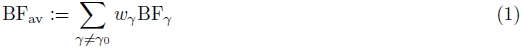

where the weights *w*_*γ*_ can be chosen to reflect the relative prior plausibility of different values of *γ*. Here we use the default weights suggested in [22], which come from putting uniform prior probability on the number of associated phenotypes *A* = |*D* ∪ *I*| = 1,…, *d* and, conditional on *A*, uniform prior probability on |*D*| = 1,…, *A*.

In cases where there is strong evidence against the null hypothesis, these methods can be used to compare the evidence for different alternative hypotheses. In particular, we summarize the evidence for *Y*_*j*_ being associated with *g* using the posterior probability of association P(*Y*_*j*_ ∈*D* or *I*). (In figures presented here we give these posterior probabilities conditional on *γ* ≠ *γ*_0_.)

In comparisons below, we compare the multivariate Bayes Factor BF_av_ with the Bayes Factor from the *d* univariate tests 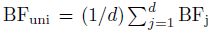 where BF_j_ denotes the Bayes Factor from a univariate association test of *Y*_*j*_. Note that BF_j_ is highly correlated with the usual *p* value testing for association between *Y*_*j*_ and *g* [23], and BF_uni_ is highly correlated with the minimum of these *p* values across the *d* phenotypes [22]. For more details on the connections between these Bayes Factors and standard tests see [22].

Calculation of the Bayes Factors requires specification of a single hyperparameters, *σ*_*a*_, which controls how large the expected genetic effects are for truly associated SNPs. In all the Bayes Factor calculations used here we averaged over *σ*_*a*_ = 0.05, 0.1, 0.2, 0.4. (See [24] for discussion.)

### Initial filtering to reduce computation of genome-wide analysis

Because the total number of possible values for *γ* in (1) is large, computing this value genome-wide is computationally intensive (e.g., 30 min/SNP for 12 phenotypes). To reduce computation we performed an initial filtering step in which, for all SNPs (7,836,525), we compute BF_all_, which is the BF for *γ* where all variables are in *D*, and BF_uni_. (Note that BF_all_ corresponds to a standard multivariate test of association, such as MANOVA; see [22].) We then performed the full analysis only on SNPs (24,601 for *Ã* and 25,175 for 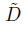) that appear promising based on this filter (log_10_ BF_uni_ > 1.3 or log_10_ BF_all_ > 1.3) The supplementary materials “all summary baseline.txt” and “all summary statin response.txt” show detailed results from the full analysis.

### Identifying deviations from multivariate normality

To identify individuals with phenotypes that appear to deviate from multivariate normality we performed multivariate outlier detection based the Mahalanobis distance as follows. Given *d*-dimensional vector phenotype vectors *y*_*1*_,…, *y*_*n*_, the Mahalanobis distance of *y*_*i*_ from the mean 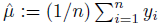 is computed using

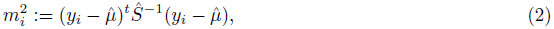

where 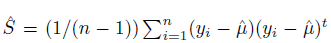 is the sample covariance matrix. For each individual we obtain a *p* value testing whether the individual’s phenotype is consistent with the multivariate normal model by using the fact that, under a multivariate normal distribution, 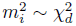.

## 3 Results

### The effect of statin treatment on LDL subfractions

Figure 1 shows the mean and standard deviation of the absolute and fractional change in LDL subfractions, by study/cohort. The three cohorts show similar trends, with stronger effects in CAP than PRINCE, presumably reflecting both the different statins and doses used, and the different enrollment criteria. In particular, all three studies show a consistent trend of larger fractional change in the larger (lower-density) particles, and lower fractional change in the smaller (higher-density) particles. This is consistent with studies showing that very small LDL have low affinity for the LDL receptor [25].

**Figure 1.**
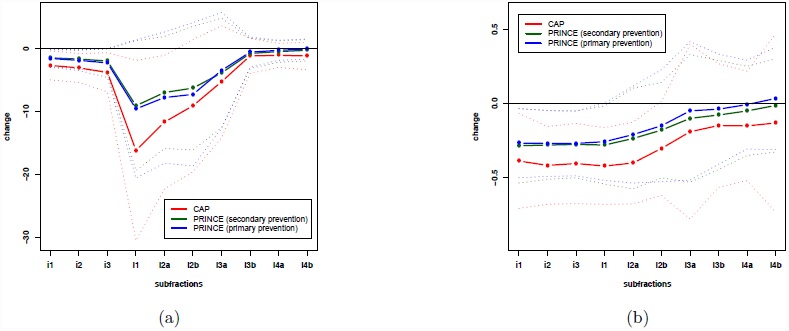
Effects of statin on cholesterol concentrations in LDL subfractions. Dashed lines show (a) mean absolute change, and (b) mean fractional change, in each study/cohort in each subfraction (see Table 1 for abbrevations); dotted lines showone ± standard deviation; subfractions are in order from lower (left) to higher (right) density lipoprotein.

### Genome-wide association analyses, irrespective of statin exposure

To identify genetic variants that are associated with LDL-C subfractions, irrespective of statin exposure, we conducted a multivariate genome-wide scan of an average of the pre-treatment measurements (*P*) and post-treatment measurements (*T*) for all twelve LDL-C related phenotypes simultaneously (10 subfractions plus Ld, and total LDL-C; Figure S1 in Supporting Information shows correlations among the phenotypes). The normalization and data processing steps used to derive this phenotype (*Ã*) are described in the methods section. The rationale for averaging pre-treatment and post-treatment measures is that it reduces the influence of environmental and temporal fluctuations in phenotype, and of measurement errors (see also [6]).

Our analysis summarizes the evidence for each SNP being associated with the multivariate phenotype data (against the null of no association) by a Bayes Factor, BF_av_, which compares the likelihood of the data under a range of alternative models, with the likelihood under the null model (see methods). Although there is no direct correspondence between Bayes Factors and *p* values, typically log_10_ BF_av_ ≈ 5 – 6 corresponds roughly to conventional levels of genome-wide significance.

We found SNPs in or near four genes (SORT1, CETP, APOE, and LPA; Figure S2 in Supporting Information) that showed very strong association signals (log_10_ BF_av_ > 15), and no other SNPs had log_10_ BF_av_ > 5. The association results, including details of the most strongly associated SNPs in each gene, are summarized in Table 2. Notably, all these signals were *much* stronger (15+ orders of magnitude) than in a univariate analysis of LDL-C alone, indicating the potential gains from using both more detailed phenotypic measurements (vs LDL-C) and multivariate analysis methods (vs univariate analysis), an issue we return to below.

**Table 2.**
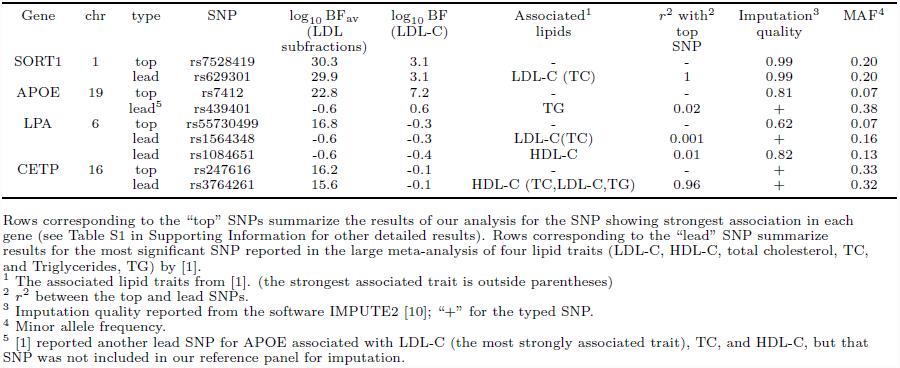
Summary of the strongest associations with LDL subfractions in our analysis.

The multivariate association analysis methods we use are based on an assumption of multivariate normality (formally, of the residuals, although when effect sizes are small this is similar to multivariate normality of the phenotypes). Although we transformed each phenotype to be univariate normal, this does not guarantee that the phenotypes will be, jointly, multivariate normal. Therefore, to check for sensitivity to deviations from multivariate normality, we repeated the analysis after removing 89 individuals who were identified as potential outliers from multivariate normality on the basis of their Mahalanobis distance from the mean (*p* < 0.01 in a test for multivariate normality; see Methods). All four of the strongest associations remained strong in this analysis; the biggest drop in signal was at rs247616 in CETP, which dropped from 16.2 to 12.2.

All these genes have been previously identified as containing SNPs associated with LDL-C in a large (univariate) meta-analysis of over 100,000 individuals [1], and also identified as being associated with lipid subfractions measured by an NMR-based method, which distinguishes three subfractions of IDLs and LDLs [2]. However, our more detailed measurements, which distinguish 10 subfractions of IDLs and LDLs, provide greater resolution to examine effects of these genetic variants on subfractions of IDLs and LDLs than previous studies. Figure 2 shows the mean value of each (normalized) subfraction phenotype by genotype class, for the most strongly associated SNP in each gene (“top SNP”). The four SNPs exhibit distinctively different patterns, presumably reflecting different roles these genes play in lipoprotein metabolism. Among them, only rs7528419 in SORT1 has an effect that is consistent in direction among all subfractions, although it primarily affects the higher-density LDL subfractions (l3b,l4a and l4b; see Table 1 for abbreviations) as reported previously [21]. This SNP is in high LD (*r*^2^ = 1) with the SNP rs12740374 that has been identified as a causal variant contributing to change in plasma LDL-C [26]. The SNP rs247616 in CETP shows a complex pattern of association, with the C allele showing an increase in both the lowest density (i1 and i2) and highest density subfractions (l3a and l3b), but a decrease in the middle of the range (l1 and l2a). Similarly, SNP rs7412 in APOE also affects a wide range of subfractions, with the minor allele increasing some (i1) and decreasing others (l1,l2a,l2b,l3a and l3b). The effect of SNP rs55730499 in LPA is concentrated entirely on the lowest density subfraction (i1) and this SNP is in high LD (*r*^2^ = 0.9) with the SNP rs10455872 that has been reported as associated with plasma lipoprotein(a) [Lp(a)] levels [27, 28]. This result probably reflects the fact that Lp(a) particles are in the same size range as the IDL1 subfraction, and likely contribute to this subfraction measurement; that is, this result may reflect an association of this SNP with Lp(a), rather than with the IDL1 subfraction [29].

**Figure 2.**
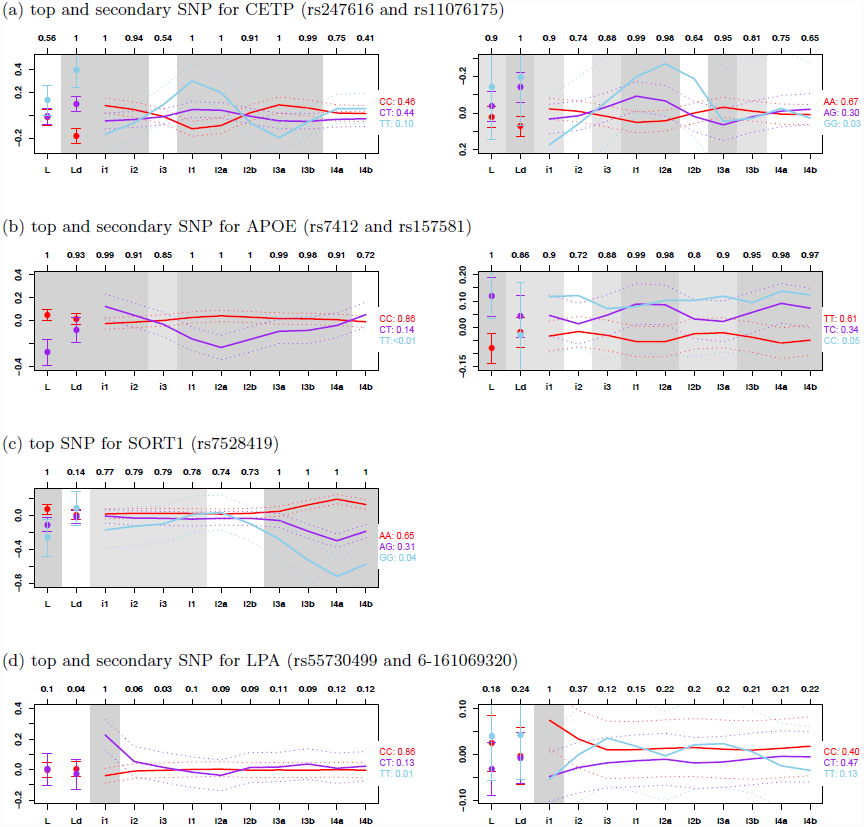
Effects of associated SNPs on LDL subfractions. Solid lines show mean (normalized) phenotype for each subfraction (see Table 1 for abbrevations) by genotype class (reference homozygotes: red, heterozygotes: purple, non-reference homozygotes: sky blue; the proportion of individuals is shown next to the genotype; sky blue lines are omitted if the proportion is ≤ 0.01); dotted lines show ±2 standard errors. Results for secondary SNPs are based on residuals from regressing out top SNPs. Grey shading indicates posterior probability of association (either directly or indirectly) for each phenotype (> 0.9: dark grey, > 0.75: light grey, < 0.75: white; raw numbers given at top of figure). Note that, because the minor alleles have opposite effects on total HDL-C at the two SNPs in CETP, the *y*-axis is reversed for the secondary SNP to emphasize the similar shapes of the curves.

### Multiple Independent Associations in CETP, APOE and LPA

Previous analyses (e.g. [1] and [3]) have indicated that many lipid-affecting genes contain multiple variants that independently affect lipid-related phenotypes. In addition, for example, [3] has shown that multiple variants in some genes affect phenotypes similarly (or dissimilarly). We therefore searched for additional independently-associated SNPs near these four genes (±150kb). We did this for each gene by performing a multivariate association analysis on the residuals obtained by regressing out the effects of the most strongly associated SNP in that gene. Although no secondary SNPs showed signals that would be considered “genome-wide” significant, CETP, APOE and LPA all contained secondary SNPs with moderately strong signals (log_10_ BF_av_ > 2; see Table 3 for details of the secondary SNPs). Furthermore, the secondary SNPs at CETP (rs11076175) and LPA (6-161069320) showed effects on the subfraction profiles that were, qualitatively, strikingly similar to those of the top SNP in those genes (Figure 2).

**Table 3.** Summary of the secondary associations with LDL subfractions in our analysis.

| Gene | chr | SNP | $\log_{10} \text{BF}_{\text{av}}(\text{LDL subfractions})^1$ | $\log_{10} \text{BF}(\text{LDL-C})^1$ | Imputation quality<sup>2</sup> | MAF<sup>3</sup> |
| --- | --- | --- | --- | --- | --- | --- |
| APOE | 19 | rs157581 | 2.68 | 3.62 | 0.94 | 0.22 |
| LPA | 6 | 6-161069320 | 2.65 | -0.44 | 0.61 | 0.36 |
| CETP | 16 | rs11076175 | 2.34 | -0.05 | + | 0.18 |

See Table S1 in Supporting Information for other detailed results.
<sup>1</sup> Results from a multivariate association analysis on the residuals obtained by regressing out the effects of the most strongly associated SNP in each gene.
<sup>2</sup> Imputation quality reported from the software IMPUTE2 [10]; “+” for the typed SNP.
<sup>3</sup> Minor allele frequency.

In contrast the secondary SNP at APOE (rs157581) showed a very different pattern to the top SNP, with the minor allele showing a consistent increase in all subfractions. That is, the top and secondary SNPs in CETP (and also LPA) have similar phenotypic consequences, possibly reflecting a similar underlying molecular effect, whereas the two SNPs in APOE have different phenotypic consequences, presumably reflecting different underlying molecular effects. The top SNP (rs7412) for APOE is one of two nonsynonymous SNPs in APOE exon 4 that define the *ε*_1_/*ε*_2_/*ε*_3_ haplotype system. The other variant (rs429358) has a modest association in the secondary analysis (log_10_ BF_av_ > 1.77) and is not in LD with the top SNP (*r*^2^ < 0.01).

The CETP locus is of considerable interest in light of controversies surrounding the relation of CETP genetic variation to risk of cardiovascular disease, and the failure of pharmacologic inhibitors of CETP to reduce this risk in recent clinical trials [30]. Given the very striking patterns of association at CETP, we sought to replicate this association in an independent population. To this end, we obtained LDL subfraction measurements, measured by ion mobility technology, for up to 6745 individuals of European ancestry from the JUPITER study (see Methods). To replicate the results for our top SNP, we estimated its effect on each subfraction using simple linear regression; to replicate results for the secondary SNP we estimated its effect on each subfraction after controlling for the top SNP. In both cases these replication data show effect size estimates that are highly concordant with those in our original study population (Figure 3), demonstrating that the highly significant initial findings also generalize to other populations.

**Figure 3.**
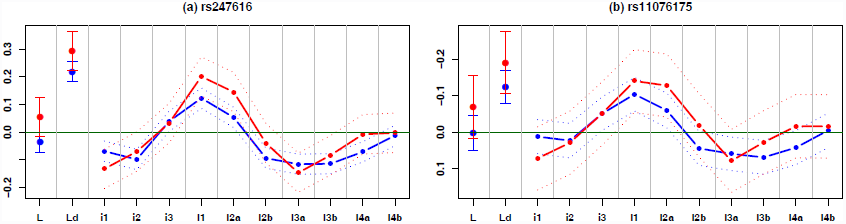
Replication of two independent associations in CETP (top SNP rs247616 and secondary SNP rs11076175). Dashed lines show estimated effect sizes on (normalized) phenotype for each subfraction in our study (red) and JUPITER (blue); dotted lines show2 standard errors. Note that, the y-axis is reversed for the secondary SNP as in Figure 2.

As CETP has been known for strong effect on total HDL-C (the top and secondary SNPs have log_10_ BF > 20 in our study), we investigated whether the association of the top and secondary SNPs with subfractions is partially mediated by association with HDL-C. To do this, for the five phenotypes, ‘Ld’, ‘i1’, ‘l1’, ‘l2a’, and ‘l3a’, with strong association signals without controlling for HDL-C (p-value < 0.005 from a standard simple linear regression), we examined how effect on the subfractions changes after controlling for HDL-C. The effect size of each SNP after controlling HDL-C was obtained by including the SNP and HDL-C as a covariate in a standard multiple linear regression. The percentage changes in effect size of the top SNP rs247616 after controlling for HDL-C are -72.7, 4.4, -89.6, -67.4, and -97.5 for ‘Ld’, ‘i1’, ‘l1’, ‘l2a’, and ‘l3a’, respectively (similar changes for the secondary SNP rs11076175); see Figure S3 in Supporting Information. Thus effect size for ‘i1’ is essentially unaffected by controlling for HDL-C, whereas effects on other subfractions are very substantially reduced (after controlling for HDL-C, the signal for the top SNP in analyses of 12 subfractions dropped from 16.2 to 3.43). These results are consistent with the SNP’s effect on LDL subfractions being largely mediated through its effect on HDL-C.

### Comparisons with previous studies

Ref [2] performed association studies on lipoprotein subfraction measurements obtained by NMR-based methods. Ref [3] used an NMR-based serum metabolomics platform to quantify 216 metabolic variables, some of which could be related to the subfraction measurements used here. For example, large LDL and small LDL subfractions measured by an NMR-based method correspond most closely to LDL1/LDL2a and LDL3a/LDL3b, respectively, in our study (see Table 2 in [31]).

Compared with these previous studies, the association results we report here for the top SNP in CETP are mostly novel. Our top SNP in CETP (rs247616) is not in high LD (*r*^2^ < 0.5) with any SNP reported as associated with NMR-based subfractions in [2]. The SNP is in high LD with a SNP (rs3764261) reported by [3], but they found no significant association between this SNP and any LDL-related measurements (their Table 1, bottom line).

The secondary SNP we report for CETP (rs11076175) is in high LD (*r*^2^ = 0.95) with one (rs7499892) of the SNPs reported in [2]. Although [2] makes no mention of the distinctive patterns of association at this SNP, the estimated effects reported in their supplementary information (Figure S4 in [2]) shows patterns concordant with those we highlight here: the effects on total IDL and small (i.e. high density) LDL particles are opposite in direction to the effect on large (low-density) LDL particles. Indeed, further examining their Figure S4 we note that there are other SNPs in CETP, not in high LD (*r*^2^ < 0.5) with either the top or secondary SNP in our analysis, that show similar distinctive association patterns, lending further support to the idea that many of the SNPs affecting lipids in CETP may have similar affects on subfractions, perhaps through a shared mechanism.

At APOE, both our top and secondary associations are distinct from those reported in [2] (*r*^2^ < 0.5 with any association reported there). Our top SNP (rs7412) was reported in the serum metabolomics study [3], with their most strongly associated phenotype being ‘enzymatically measured LDL-C’ (Table 2 and Table S10 in [3]).

The top SNP in SORT1 from our analysis is in high LD (*r*^2^ = 1) with SNPs reported as associated in the previous NMR-based studies (rs646776 in [2] and rs629301 in [3]). However, neither of these previous studies has the resolution to highlight the distinctive patterns of effects highlighted in the gradient gel electrophoresis results, with by far the strongest associations being with the smallest and densest LDL subfractions (3b/4a/4b).

The two independent associations for LPA from our analysis were not reported in either of [2, 3]. Since these association may be due to associations with Lp(a), rather than with an LDL subfraction, this difference may reflect different sensitivities of the technologies used to Lp(a). Interestingly, our secondary SNP (6-161069320) is not in LD (*r*^2^ < 0.25) with any of five SNPs that have been reported in the NHGRI GWAS catalog [32] as associated with Lp(a) levels, and so may represent a novel association with Lp(a).

### Genome-wide association analysis of statin response

To identify SNPs associated with response to statin we performed a multivariate genome-wide association analysis using the difference between post-treatment and pre-treatment measures of each phenotype as a multivariate outcome. These differences, 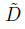, were obtained using the standardized (quantile-normalized) phenotypes (see Methods), so this association analysis targets genetic variants whose *standardized* effect on phenotype changes pre- and post- treatment. For example, a SNP that explains 1% of phenotypic variance before treatment and 2% after treatment would be associated with response to statin in our analysis.

Only one SNP (rs6430626) had log_10_ BF_av_ > 5 (actual log_10_ BF_av_ = 5.3). The phenotypes contributing to this strong multivariate signal included a modest (univariate) association with total LDL-C response (log_10_ BF = 1.7). However, imputation quality at this SNP was low (0.25), and a substantial part of the association signal appeared to be driven by deviations from multivariate normality: on removing 96 individuals detected as having outlying (multivariate) phenotypes at the 0.01 significance threshold (see Methods), log_10_ BF_av_ fell to 2.3. Furthermore, neither this SNP, nor other SNPs in LD with it, showed even nominal associations with (total) LDL response to statins in the JUPITER study (*p* values near 0.9).

Although no other SNPs showed strong evidence for association with statin response in our analysis, our data also provide the potential opportunity to assess which subfractions of LDL are most likely responsible for previously-observed associations with LDL-C response to statin in the much larger JUPITER study [7]. To this end we examined the three SNPs (near the genes ABCG2, APOE and LPA) highlighted in [7] as being associated with fractional change in LDL-C (at *p* value < 5 × 10^−8^).

The SNP rs1481012 near ABCG2 showed no signal for association with response to statin in either the subfractions or total LDL-C (log_10_ BF_av_ = –0.65). Possibly this lack of signal could reflect our small sample size, although it could also be due to the different statin drugs used (CAP used simvastatin; PRINCE used pravastatin; JUPITER used rosuvastatin).

The SNPs rs7412 in APOE and rs10455872 in LPA had modest association signals with statin response in our data (rs7412: log_10_ BF_av_ = 1.6 and rs10455872: log_10_ BF_av_ = 0.8). Both these SNPs are associated with LDL phenotypes both before and after statin treatment, so these associations with statin response must reflect a difference in the *strengths* of these associations in the untreated vs treated condition. That is, they show a statistical interaction with treatment. To gain additional insights into these interactions we examined whether the effect sizes are stronger in the treated or the untreated data. The APOE SNP showed consistently stronger estimated effects in the treated condition compared with the untreated condition, across the smaller particles (l2a-l4a; Figure 4a). The LPA SNP also showed stronger effects in treated vs untreated conditions, but with the effects concentrated in the IDL1, and, to a lesser extent, IDL2 subfraction (Figure 4b). As noted above, the association of this SNP with the IDL1 subfraction likely reflects, at least in part, the effect of this SNP on Lp(a) particles. The stronger association with IDL1 post-treatment may therefore be explained by the fact that Lp(a) levels are resistant to statin treatment [27], and so will contribute more strongly to the post–statin measures than the pre-statin measures.

**Figure 4.**
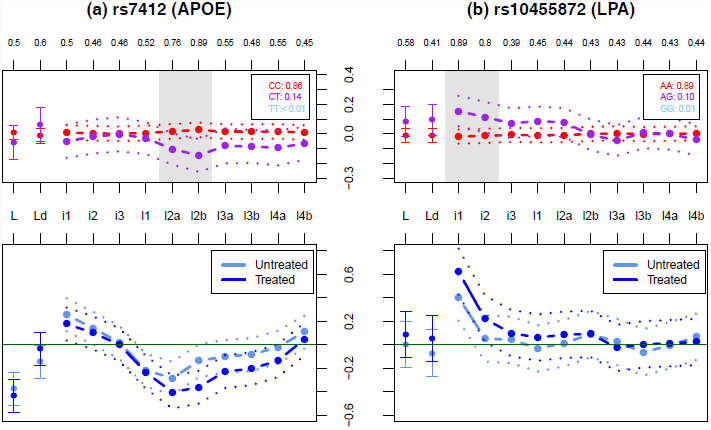
Effects of associated SNPs (rs7412 in APOE and rs10455872 in LPA) on treated and untreated measures of LDL subfractions, and on their difference. Upper panels show the effects of the SNPs on the difference between treated and untreated measures of LDL subfractions. Labels and colors are as in Figure 2. Lower panels show estimated effect size (dashed lines) and ±2 standard errors (dotted lines) in the treated and untreated (normalized) phenotype for each subfraction.

### Comparison of multivariate and univariate association analyses

In our genome-wide association analysis irrespective of statin exposure, all four of the strongest associations (in or near CETP, APOE, SORT1, and LPA) showed very much stronger association signals in the multivariate analysis than in a univariate analysis of LDL-C. Two factors could contribute to this increased signal: first, our use of more-detailed phenotypes (the subfractions), any of which could show individually stronger association signals than total LDL-C; and second, our analysis of these phenotypes simultaneously, in a multivariate way, rather than one at a time. We now investigate the contributions of these two factors to increased association signals in more detail. To do this we compare three different Bayes Factors that test for association: the BF based on univariate analysis of LDL-C (BF_ldl_), the BF based on univariate analysis of all 12 phenotypes (BF_uni_) and the BF based on multivariate analysis of all 12 phenotypes (BF_av_). We observed above that for the strongest associations BF_av_ was many orders of magnitude larger than BF_ldl_. We now attempt to decompose this gain in signal into two parts, using the identity

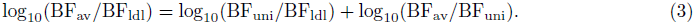

Intuitively, we can think of the first term as capturing the gain (or loss) from the more detailed subfraction measurements, and the second term as capturing the gain (or loss) in going from a univariate to a multivariate analysis.

To increase the number of SNPs contributing to this assessment, we compare not only the very strongest associations, but also more modest association signals. To attempt to ensure that these more modest associations nonetheless reflect likely genuine associations we focus on SNPs in or near the 95 genes (±150kb) identified as being associated with lipid traits in [1] (“Global Lipids genes”). We included all the genes from this study, and not only those associated with LDL-C, because we wanted to allow for the fact that, as illustrated in our main analysis, SNPs may be strongly associated with LDL subfractions without being strongly associated with LDL-C. We include in our comparisons any SNP for which any of the BFs (BF_av_, BF_ldl_, BF_uni_) exceed 10^3^. Given the modest nature of some of these associations we cannot be completely confident that all of them are genuine, although it seems reasonable to believe that most of them are.

Figure 5 breaks down the gain (or loss) from using BF_av_ vs BF_ldl_ (panel a) into a component due to the more detailed measurements (panel b) and a component due to the multivariate analysis (panel c). For the majority of associations, both the use of more detailed phenotypic measurements and the use of multivariate association analysis contribute to an increase in the association signal. For some associations (e.g. in SORT1, APOA1) the main gain comes from the more detailed phenotype measurements; for others (e.g. some associations in APOE, LPA) the increase in association signal is greater in moving from the univariate to multivariate analysis.

**Figure 5.**
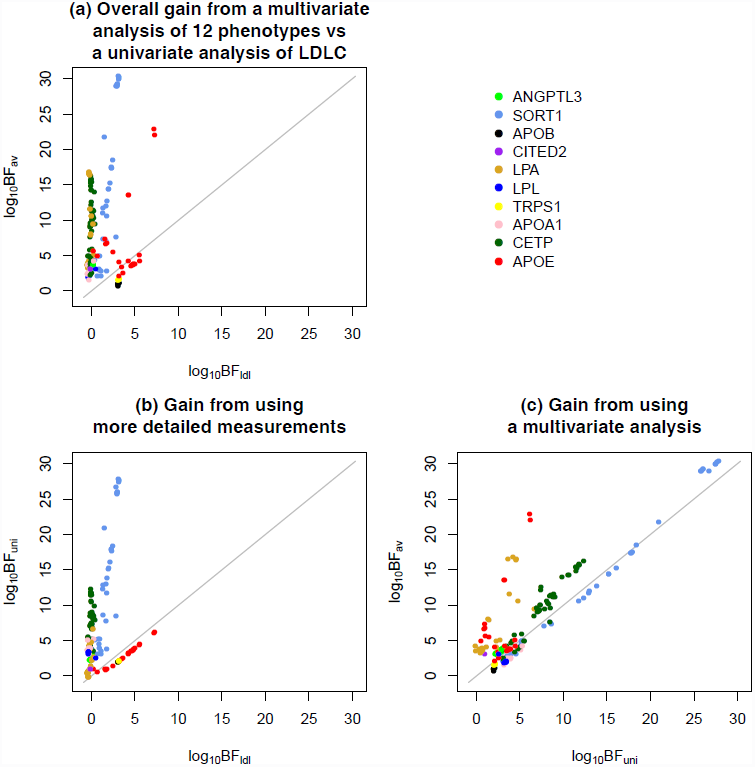
Decomposition of the gain (or loss) from a multivariate analysis of 12 phenotypes vs a univariate analysis of LDL-C into two components: one from using more detailed measurements, and one from using a multivariate analysis. Plotted are (a) log_10_ BF_av_ (the BF based on multivariate analysis of all 12 phenotypes) vs log_10_ BF_ldl_ (the BF based on univariate analysis of LDL-C), (b) log_10_ BF_uni_ (the BF based on univariate analysis of all 12 phenotypes) vs log_10_ BF_ldl_, and (c) log_10_ BF_av_ vs log_10_ BF_uni_. SNPs are colored according to the nearest gene.

It is natural to ask what features of the different associations explain these different behaviors, and, more generally, under what circumstances more detailed measurements or multivariate analyses may prove most helpful. For multivariate analyses, the general answer is that the gains in multivariate analyses are largest when the effects are discordant with the correlation among phenotypes. For example, for two positively correlated phenotypes, the gains of multivariate analysis will be greatest when the effects on the two phenotypes have opposite signs; there will also be substantial gains when one phenotype has an effect and the other has zero effect; the gains will be smallest when both phenotypes have an effect of the same sign. See [22] for more discussion. This observation helps explain the gain of the multivariate analyses for APOE and LPA associations, both of which exhibit some discordance between effect and correlation structure: for APOE, some negatively-correlated subfractions (e.g. l1 and l3a; see Figure S1 in Supporting Information) show effects in the same direction (Figure 2); for LPA, subfraction i1 shows a substantive effect, and many subfractions correlated with this subfraction (both positively and negatively) show no effect. It is harder to generalize as to when more detailed measurements will increase power to detect associations. In our study, the largest gains in power from the subfraction measurements were at SORT1 and at CETP, and these gains appear to be for different reasons. At CETP, the gain is due to the genetic variants affecting subfractions having effects in different directions, which approximately “cancel out” so that the effect on total LDL-C is small compared with the effects on individual subfractions. At SORT1, the gain is because the genetic variants affect primarily the smaller (l3b,l4a,l4b) subfractions, which together represent only a small proportion of total LDL-C, and contribute correspondingly little to overall variation in total LDL-C (See Table 1). That is, the genetic effects on these subfractions are diluted by variation in the other more abundant subfractions (l1-l3a) when testing for association with LDL-C.

We note that, of course, power is only one potential benefit of the subfraction measurements: even when the more detailed measurements do not increase power to detect associations, being able to estimate the effects on different subfractions may still be useful (e.g. because the different subfractions have different biologic and clinical implications).

## Discussion

We have presented an association analysis of genetic variants with detailed measurements of IDL/LDL subfractions, and highlighted several SNPs that, while only weakly associated with total LDL-C in our study, are strongly associated with the subfractions. Perhaps the most interesting of the associations are the two independently-associated SNPs in CETP. From previous studies these variants are known to be very strongly associated with plasma HDL-C (log_10_ BF > 20 in our study), and less strongly associated with total LDL-C. For example, the lead SNP in [1] explains 1.5% of the variance in HDL-C, but only 0.06% of the variance in LDL-C (see Supporting Information). Our data demonstrate that, nonetheless, these variants are very strongly associated with multiple subfractions of LDL, with effects in different directions on different subfractions approximately cancelling each other out to yield a small overall effect on total LDL-C. Interestingly, these independent variants have similar effects on the different subfractions, consistent with a shared underlying functional mechanism that may involve selective CETP-mediated cholesterol-triglyceride exchange between HDL and specific subclasses of apoB-containing particles [33]. The findings complement previous analyses of LDL subfractions in two hyperalphalipoproteinemic patients with genetic deficiency of CETP [34], and recent data on the effects of the CETP inhibitor anacetrapib on lipoprotein subfraction concentrations [33], both of which indicate that CETP influences LDL subclasses in a manner consistent with the CETP genotype associations shown here. Specifically, reduced CETP activity in both instances results in an increase in small LDL particles together with a reduction in mid-sized LDL. While the mechanism for this effect is not known, it has been suggested that it is related to the existence of separate pathways giving rise to larger vs. mid-sized and smaller LDL [18, 34], with CETP deficiency selectively resulting in triglyceride enrichment of particles in the second of these pathways, and a resultant shift from mid-sized to smaller LDL through the action of lipase activity [33,34].

Our study also provides a practical illustration of the potential benefits of joint association analysis of multiple related phenotypes in genetic association studies, rather than considering each phenotype one at a time. While in principle the potential benefits of joint analysis are well established, and have been highlighted in several recent papers (e.g. [35–42]), in practice the vast majority of association studies of multiple phenotypes continue to rely on univariate analyses. We hope that our comparisons provide further motivation to investigators to consider joint multivariate analyses as a tool to help improve power to detect genetic associations.

## Acknowledgments

We thank Mathew Barber for help in the early stages of this work; Bryan Howie for help with imputation; and Ellen Leffler for helpful comments on an earlier version of the manuscript. We thank the members of the Pritchard, Przeworski, and Stephens labs for helpful discussions.

## Supporting Information

### Computing proportion of variance in phenotype explained by a given SNP (PVE)

[1] provides sample size, minor allele frequency (MAF), effect size, and standard error of effect size for each reported SNP (see Supplementary Table 2 in [1]). We estimated PVE using the information from [1] as follows. Variance in phenotype (*Y*) can be decomposed into two components:

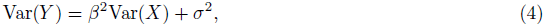

where *β* is effect size of genetic variant (*X*). The first component (*β*^2^Var(*X*)) captures variance explained by the genetic variant *X* and the second component (*σ*^2^) captures the remaining variance that can be explained by environmental factors or other genetic variants. We can estimate *β*^2^Var(*X*) by 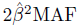 (1-MAF), where 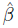 and MAF are effect size estimate and minor allele frequency for the genetic variant *X*, respectively. From a simple linear regression model (X and Y as covariate and response),

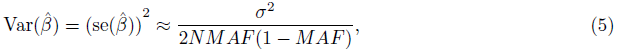

where *N* is sample size and se(*β*^ˆ^) is standard error of effect size for the genetic variant *X*. Therefore,

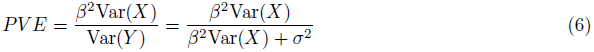

can be estimated by

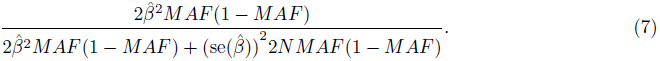

We compute PVE for HDL-C (LDL-C) by using the information for the most strongly associated SNP rs3764261 (rs247616). Note that rs3764261 is in high LD (*r*^2^ = 0.96) with rs247616.

**Figure S1: Spearman correlations among 16 phenotypes**. In addition to 12 phenotypes considered in our main analysis, correlations with triglycerides (Tg), HDL-cholesterol (H), total cholesterol (T), ApoB levels (A) are also included. See Table 1 in the main text for abbrevations for 12 phenotypes. Correlations are computed by using (a) (normalized) pre-treatment measurements (*P* in the methods section); (b) (normalized) post-treatment measurements (*T* in the methods); (c) (normalized) averages of pre-treatment measurements and post-treatment measurements (*Ã* in the methods); (d) (normalized) differences between post-treatment and pre-treatment measures (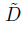 in the methods).

**Figure S2: Regional plots of the SORT1, APOE, LPA and CETP**. Regional plots of the four loci are created by using the tool LocusZoom [43] (with log_10_ BF_av_ in y-axis). Purple diamond is the top SNP with the strongest association in each locus. Each circle corresponds to a SNP whose color indicates the linkage disequilibrium with the top SNP. LDL subfraction associations around (a) SORT1, (b) APOE, (c) LPA, and (d) CETP.

**Figure S3: Effect size with ±2 standard error before (blue) and after (red) controlling for HDL-C for each subfraction**. (a) The top SNP rs247616 and (b) the secondary SNP rs11076175 in CETP. Note that, because the minor alleles have opposite effects on total HDL-C at the two SNPs, the y-axis is reversed for the secondary SNP to emphasize the similar pattern of effect size.

**Table S1.** Summary of the top and secondary associations with LDL subfractions in our analysis.

| Gene | SORT1 | APOE | APOE | LPA | LPA | CETP | CETP |
| --- | --- | --- | --- | --- | --- | --- | --- |
| chr | 1 | 19 | 19 | 6 | 6 | 16 | 16 |
| type | top | top | secondary<sup>1</sup> | top | secondary | top | secondary |
| SNP | rs7528419 | rs7412 | rs157581 | rs55730499 | 6-161069320 | rs247616 | rs11076175 |
| LDL-C (L) |  |  |  |  |  |  |  |
| log<sub>10</sub> BF<sup>2</sup> | 3.07 | 7.18 | 3.62 | -0.28 | -0.44 | -0.12 | -0.05 |
| PP of direct association<sup>3</sup> | 0.95 | 0 | 0.96 | 0.07 | 0.06 | 0.15 | 0.27 |
| PP of indirect association<sup>3</sup> | 0.05 | 1 | 0.04 | 0.03 | 0.12 | 0.41 | 0.63 |
| PP of no association<sup>3</sup> | 0 | 0 | 0 | 0.9 | 0.82 | 0.44 | 0.1 |
| LDL peak particle diameter (Ld) |  |  |  |  |  |  |  |
| log<sub>10</sub> BF | -0.48 | -0.08 | -0.36 | -0.28 | -0.41 | 13.39 | 3.24 |
| PP of direct association | 0 | 0.7 | 0.41 | 0.01 | 0.07 | 0 | 0.98 |
| PP of indirect association | 0.13 | 0.23 | 0.45 | 0.03 | 0.17 | 1 | 0.02 |
| PP of no association | 0.86 | 0.07 | 0.14 | 0.96 | 0.76 | 0 | 0 |
| IDL1 (i1) |  |  |  |  |  |  |  |
| log<sub>10</sub> BF | -0.20 | 1.47 | 0.2 | 5.25 | 1.17 | 2.06 | -0.01 |
| PP of direct association | 0.74 | 0.01 | 0.55 | 0 | 1 | 0.6 | 0.31 |
| PP of indirect association | 0.03 | 0.98 | 0.35 | 1 | 0 | 0.4 | 0.58 |
| PP of no association | 0.23 | 0.01 | 0.1 | 0 | 0 | 0 | 0.1 |
| IDL2 (i2) |  |  |  |  |  |  |  |
| log<sub>10</sub> BF | -0.06 | 0.22 | -0.25 | -0.09 | -0.26 | 0.16 | -0.39 |
| PP of direct association | 0.76 | 0.22 | 0.61 | 0 | 0.15 | 0.65 | 0.27 |
| PP of indirect association | 0.03 | 0.7 | 0.12 | 0.06 | 0.21 | 0.29 | 0.47 |
| PP of no association | 0.21 | 0.09 | 0.28 | 0.94 | 0.63 | 0.06 | 0.26 |
| IDL3 (i3) |  |  |  |  |  |  |  |
| log<sub>10</sub> BF | -0.12 | -0.32 | 0.01 | -0.24 | -0.46 | -0.38 | -0.24 |
| PP of direct association | 0.76 | 0.61 | 0.45 | 0 | 0.11 | 0.31 | 0.19 |
| PP of indirect association | 0.03 | 0.24 | 0.43 | 0.03 | 0.01 | 0.23 | 0.69 |
| PP of no association | 0.21 | 0.15 | 0.12 | 0.97 | 0.88 | 0.46 | 0.12 |
| LDL1 (l1) |  |  |  |  |  |  |  |
| log<sub>10</sub> BF | -0.31 | 1.69 | 0.97 | -0.29 | -0.46 | 5.75 | 1.42 |
| PP of direct association | 0.75 | 0.71 | 0.45 | 0.07 | 0.07 | 0.86 | 0.21 |
| PP of indirect association | 0.03 | 0.29 | 0.54 | 0.03 | 0.08 | 0.14 | 0.78 |
| PP of no association | 0.22 | 0 | 0.01 | 0.9 | 0.85 | 0 | 0.01 |
| LDL2a (l2a) |  |  |  |  |  |  |  |
| log<sub>10</sub> BF | -0.41 | 5.02 | 1.08 | -0.29 | -0.43 | 2.60 | 1.06 |
| PP of direct association | 0.71 | 0.74 | 0.39 | 0.06 | 0.06 | 0.86 | 0.21 |
| PP of indirect association | 0.03 | 0.26 | 0.59 | 0.03 | 0.16 | 0.14 | 0.77 |
| PP of no association | 0.26 | 0 | 0.02 | 0.91 | 0.78 | 0 | 0.02 |
| LDL2b (l2b) |  |  |  |  |  |  |  |
| log<sub>10</sub> BF | -0.12 | 2.51 | -0.08 | -0.26 | -0.45 | -0.31 | -0.43 |
| PP of direct association | 0.68 | 0.39 | 0.53 | 0.06 | 0.05 | 0.79 | 0.25 |
| PP of indirect association | 0.05 | 0.61 | 0.28 | 0.03 | 0.15 | 0.12 | 0.39 |
| PP of no association | 0.27 | 0 | 0.2 | 0.91 | 0.8 | 0.09 | 0.36 |
| LDL3a (l3a) |  |  |  |  |  |  |  |
| log<sub>10</sub> BF | 1.32 | 0.63 | -0.15 | -0.29 | -0.46 | 2.68 | 0.08 |
| PP of direct association | 0.97 | 0.68 | 0.36 | 0.08 | 0.05 | 0.81 | 0.22 |
| PP of indirect association | 0.03 | 0.31 | 0.54 | 0.03 | 0.15 | 0.19 | 0.73 |
| PP of no association | 0 | 0.01 | 0.1 | 0.89 | 0.8 | 0 | 0.05 |
| LDL3b (l3b) |  |  |  |  |  |  |  |
| log<sub>10</sub> BF | 11.84 | 0.41 | 0.30 | -0.29 | -0.45 | 0.45 | -0.40 |
| PP of direct association | 0.94 | 0.54 | 0.41 | 0.06 | 0.05 | 0.87 | 0.19 |
| PP of indirect association | 0.06 | 0.44 | 0.54 | 0.03 | 0.16 | 0.13 | 0.62 |
| PP of no association | 0 | 0.02 | 0.05 | 0.91 | 0.79 | 0.01 | 0.19 |
| LDL4a (l4a) |  |  |  |  |  |  |  |
| log<sub>10</sub> BF | 28.92 | -0.13 | 1.52 | -0.29 | -0.38 | -0.52 | -0.44 |
| PP of direct association | 0 | 0.54 | 0.63 | 0.09 | 0.06 | 0.62 | 0.21 |
| PP of indirect association | 1 | 0.37 | 0.35 | 0.03 | 0.15 | 0.13 | 0.54 |
| PP of no association | 0 | 0.09 | 0.02 | 0.88 | 0.79 | 0.25 | 0.25 |
| LDL4b (l4b) |  |  |  |  |  |  |  |
| log<sub>10</sub> BF | 12.99 | -0.14 | 0.86 | -0.28 | -0.31 | -0.53 | -0.44 |
| PP of direct association | 0.97 | 0 | 0.4 | 0.09 | 0.06 | 0.16 | 0.26 |
| PP of indirect association | 0.03 | 0.71 | 0.57 | 0.03 | 0.16 | 0.26 | 0.4 |
| PP of no association | 0 | 0.28 | 0.03 | 0.88 | 0.78 | 0.59 | 0.35 |

Abbreviations for each phenotype are in parentheses.
<sup>1</sup> Results for secondary SNPs are from analysis on the residuals obtained by regressing out the effects of the most strongly associated SNP (top SNP) in each gene.<sup>2</sup> log<sub>10</sub>Bayes Factor from a univariate analysis of each phenotype.<sup>3</sup> Marginal posterior probability of direct, indirect, no association for each phenotype that is computed from our multivariate analysis. Here, we compute these posterior probabilities conditional on at least one phenotype being directly associated.

